# Phylogenomic species tree estimation in the presence of incomplete lineage sorting and horizontal gene transfer

**DOI:** 10.1101/023168

**Authors:** Ruth Davidson, Pranjal Vachaspati, Siavash Mirarab, Tandy Warnow

## Abstract

**Background:** Species tree estimation is challenged by gene tree heterogeneity resulting from biological processes such as duplication and loss, hybridization, incomplete lineage sorting (ILS), and horizontal gene transfer (HGT).

Mathematical theory about reconstructing species trees in the presence of HGT alone or ILS alone suggests that quartet-based species tree methods (known to be statistically consistent under ILS, or under bounded amounts of HGT) might be effective techniques for estimating species trees when *both* HGT and ILS are present.

**Results:** We evaluated several publicly available coalescent-based methods and concatenation under maximum likelihood on simulated datasets with moderate ILS and varying levels of HGT. Our study shows that two quartet-based species tree estimation methods (ASTRAL-2 and weighted Quartets MaxCut) are both highly accurate, even on datasets with high rates of HGT. In contrast, although NJst and concatenation using maximum likelihood are highly accurate under low HGT, they are less robust to high HGT rates.

**Conclusion:** Our study shows that quartet-based species-tree estimation methods can be highly accurate under the presence of both HGT and ILS. The study suggests the possibility that some quartet-based methods might be statistically consistent under phylogenomic models of gene tree heterogeneity with both HGT and ILS.

## Background

A species phylogeny is a graphical model of the common evolutionary history of a group of species, and is most often represented as a phylogenetic tree or phylogenetic network [1]. A species phylogeny gives valuable information about protein functions [2, 3, 4], host-parasite relationships [5], etc.

However, species tree estimation is difficult, due to multiple biological processes, including recombination [6], duplication and loss [7], hybridization [8], incomplete lineage sorting (ILS) [9], and horizontal gene transfer (HGT) [10], that can cause a given genomic locus to have a tree that is different from the species tree. As a result, multiple loci are needed to estimate a species phylogeny with high accuracy.

Of the many sources of gene tree discord, the one that has received the greatest attention is ILS, which is modeled by the multi-species coalescent (MSC) model [11]. An MSC model tree has a rooted tree *T*, leaf-labelled by a set of species, and is given with branch lengths in coalescent units. Gene trees evolve within the species tree, in a backwards process described by the MSC; thus, lineages “coalesce” on the branches of the tree, as they move from the leaves of the species tree towards the root. When two lineages fail to coalesce on the earliest branch in which they can coalesce, this can result in a gene tree having a different topology than the species tree.

Under the MSC model, each species tree defines a probability distribution on gene trees, and the species tree can be identified uniquely from this distribution. Hence, one type of technique (called a “summary method”) for estimating species trees under the MSC operates by first estimating gene trees for a set of different loci, and then uses this estimated distribution on gene trees to estimate the species tree. A summary method is said to be statistically consistent under the MSC model if, as the number of loci and sites per locus go to infinity, the estimated species tree returned by the method will converge in probability to the true species tree [12]. Many statistically consistent summary methods have been developed for estimating species trees when gene discordance is due to ILS [13, 14, 15, 16, 17, 18, 19].

Despite advances in developing statistically consistent methods for species tree estimation that are robust to ILS, by far the most common technique for estimating a species tree is concatenation analysis, in which the sequence alignments for the different loci are combined into one large supermatrix, and then a phylogeny is estimated on the alignment using maximum likelihood [20, 21]. This type of approach, however, is sometimes not statistically consistent under the multi-species coalescent model [22, 12] in the presence of ILS. Hence, even though concatenation often has good accuracy (even under conditions with moderately high ILS levels) [23, 24, 25], a large effort has been made to develop alternative methods that are provably robust to ILS and have good accuracy on realistic conditions.

For very small datasets, Bayesian methods such as BEST [26], *BEAST [27] or BUCKy-pop [28] (the population tree from BUCKy) can provide excellent accuracy; however, these methods are too computationally intensive to use on even moderate sized datasets with hundreds to thousands of loci and 30 or more species [29, 30].

Of the currently available coalescent-based methods, ASTRAL-2 [19], MP-EST [13], and NJst [17] have emerged as the most accurate of the methods that can run on datasets with 50 or more species and hundreds to thousands of loci. However, the comparison among these methods shows that MP-EST is typically not as accurate as NJst and ASTRAL-2 and is also much slower than both [19]. Some newer statistically consistent methods have also been developed (e.g., SVDquartets [31]), but have not yet been sufficiently evaluated in terms of their accuracy and scalability in comparison to other coalescent-based methods.

Some of the most commonly used coalescent-based methods estimate species trees by encoding each gene tree as a set of quartet trees (i.e., unrooted 4-leaf trees), and then estimate the species tree from the quartet tree frequencies. The mathematical basis of this approach is the following theorem, originally proved in [32]:

### Theorem 1

*Under the multi-species coalescent model, for every model species tree T, θ) (where θ denotes the branch lengths of T in coalescent units) and for every set χ of four leaves from T, the most probable unrooted gene tree topology on χ is identical to the species tree T restricted to leafset χ.*

Interestingly, nearly the same theorem was proven under two phylogenomic models that addressed horizontal gene transfer (HGT)! When HGT is present, the evolutionary history of the species is not really treelike, but rather requires a phylogenetic network [1]. Under HGT models, a phylogenetic network consists of an underlying species tree *T* with horizontal gene transfer edges (represented by directed edges) between branches in the tree, and each locus evolves down a tree (though not necessarily the species tree) within this network. Hence, while the species evolution is not purely treelike, the gene tree evolution *is* treelike. Furthermore, for this type of reticulate phylogeny, it is reasonable to ask whether the underlying species tree *T* can be reconstructed from gene trees estimated on the different loci.

This question has been partially answered for two models of HGT. The first models HGT events between lineages using a continuous-time Poisson process [33], and is called the *stochastic HGT model*. In a stochastic HGT model, the HGT events happen between contemporaneous lineages, either uniformly at randomly or with probability that depends on the distance between the lineages (so that events are less likely if the lineages are more distantly related). The second type of model assumes that there are HGT edges between specific pairs of branches in a species tree, commonly referred to as *highways*, along which HGT events are far more likely to occur than elsewhere in the tree; this is called the *highways HGT model* [34].

The theoretical framework for estimating the underlying species tree under these two HGT models was established in [35] (for estimating rooted species trees from rooted gene trees) and in [36] (for estimating unrooted species trees from unrooted gene trees). Specifically, [36] proved theorems that under both the stochastic HGT model and highways model, but with bounded amounts of HGT per gene, the most probable quartet tree would be topologically identical to the species tree. Note that these theorems are the equivalents of Theorem 1 under the two bounded HGT models.

Some species tree estimation methods operate by computing gene trees, encoding each computed gene tree as a set of quartet trees, and determine the dominant quartet tree for every four species (i.e., the quartet tree that appears the most frequently of the three possible unrooted quartet trees). Then, these dominant quartet trees are combined using a quartet amalgamation method (e.g., Quartets Max Cut [37] or QFM [38]). This type of species tree estimation method can be statistically consistent under the MSC model, and also under these bounded HGT models – depending on the quartet amalgamation method, as we now show.

### Theorem 2

*Let M be a summary method (i.e., a method that constructs a species tree from an input set of gene trees). Suppose that M has the property that it is guaranteed to return the unique tree compatible with the dominant quartet trees defined by its input set of gene trees, whenever the dominant quartet trees are compatible. Then M is statistically consistent under the MSC model, and also under the bounded HGT models given in [36].*

*Proof* To establish statistical consistency, we only need to prove that as the number of sites per locus and the number of loci both increase, the tree returned by the method converges in probability to the species tree. As the number of sites per locus and the number of loci both increase, the dominant quartet tree converges to the most probable quartet tree on every set *χ* of four species. Under the MSC model and also under the bounded HGT models in [36], the most probable quartet tree on any set *χ* is topologically identical to the species tree. Hence, for a large enough number of loci and large enough number of sites per locus, with probability converging to 1, the input to the quartet-based methods will be a set of gene trees such that the dominant quartet trees are all compatible with the species tree. Furthermore, the species tree will be the unique such compatibility tree, and so the method will return the true species tree. □

Similarly, we can prove the following:

### Theorem 3

*ASTRAL and ASTRAL-2 are statistically consistent under the bounded HGT models of [36].*

This proof uses Theorem 1, but is essentially identical to the proofs of statistical consistency for ASTRAL and ASTRAL-2 under the MSC model [19]; see Methods for the proof of this theorem.

Very little is known about the theoretical guarantees of any species tree estimation methods under models in which both HGT and ILS can occur. In fact, to the best of our knowledge, no methods have yet been proven statistically consistent under these conditions. We also do not know much about the empirical performance of any species tree estimation methods under these conditions. As far as we know, the only simulation study to date of the impact of both ILS and HGT on the performance of species tree estimation methods is [39], which explored the performance of two coalescent-based methods, BUCKy and BEST, on data that evolved under both processes. However, both of these methods are computationally intensive, and cannot run on even moderately large datasets (e.g., BEST is slower than *BEAST, and *BEAST is too computationally intensive to use on datasets with more than about 100 loci) [30, 29].

We report on a study evaluating the accuracy of ASTRAL-2, NJst, and weighted Quartets Max Cut (wQMC) [40], as well as unpartitioned maximum likelihood concatenation analysis (CA-ML), on simulated datasets in which gene tree discord is due to both HGT and ILS. The simulation protocol evolved gene trees down 50taxon species trees under the MSC model with a moderately high level of ILS, and allowed gene trees to then evolve with six different HGT rates (see Fig. 1). HGT rate (1) has no HGT events, and HGT rates (2)-(6) have 0.08, 0.2, 0.8, 8.0, and expected HGT events per gene, respectively. Finally, sequences evolved down each gene tree under the GTR+Gamma model.

**Figure 1.**
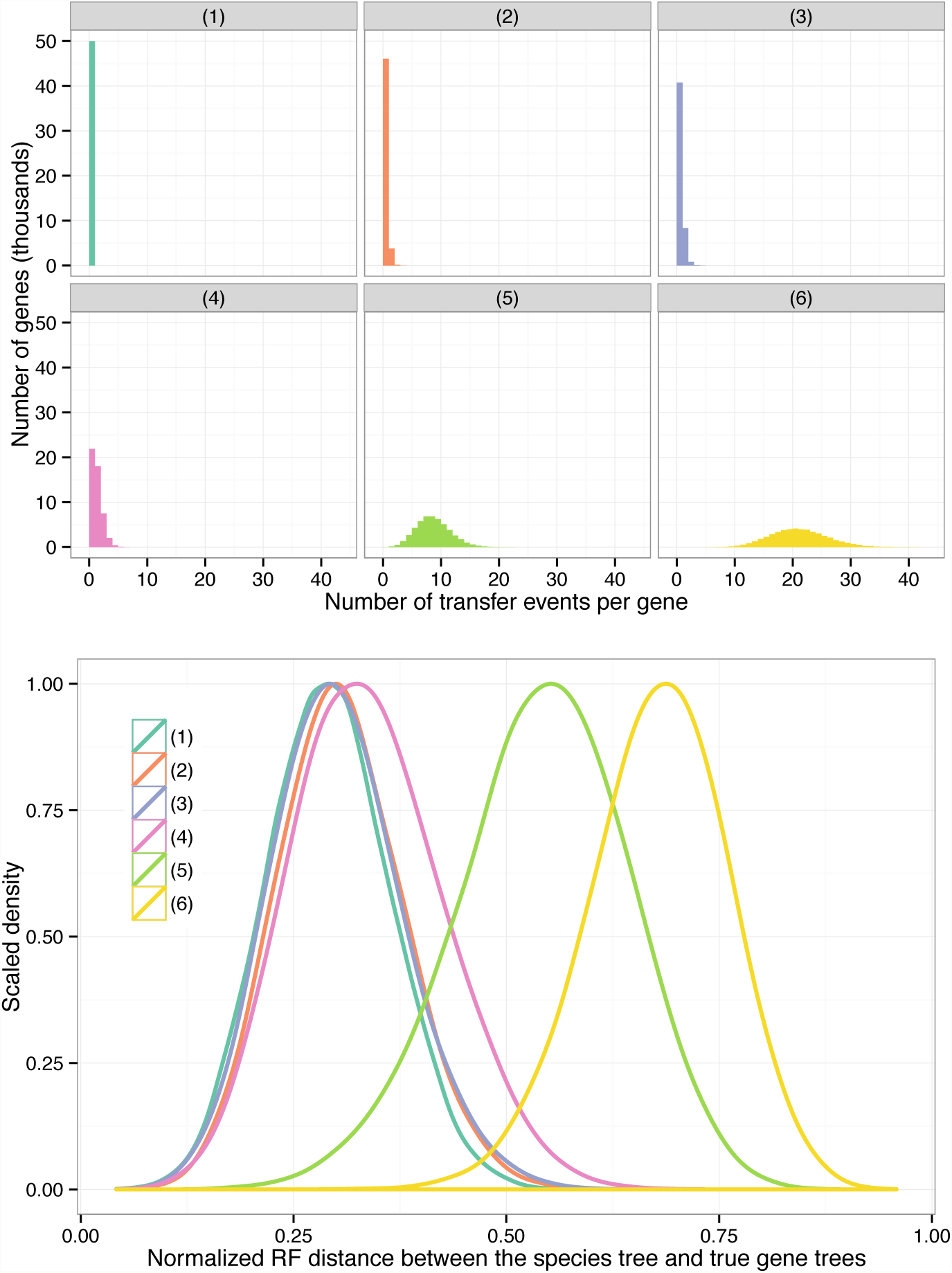
Properties of the simulated datasets. (Top) The histogram of the number of transfer events per gene across all 50,000 gene trees (50 replicates, each with 1000 genes) for all six model conditions. Note that the tree has only 51 species (50 ingroup species and one outgroup species), and therefore, model conditions (5) and (6) constitute high numbers of transfers per

We estimated gene trees on each locus using the FastTree-2 maximum likelihood software [41], and then used the summary methods on these estimated gene trees to estimate the species tree. We also concatenated the sequence alignments and ran unpartitioned FastTree-2 maximum likelihood on the concatenated superalignment. Finally, we analyzed a Cyanobacteria dataset with 11 species and 1128 genes [42], which is believed to have evolved under high levels of HGT and has been used to evaluate methods for inferring species trees in the presence of HGT [43, 40]. See Methods for additional details.

## Results

We ran 28 experiments using ASTRAL-2, NJst, wQMC, and an unpartitioned concatenated maximum likelihood analysis (CA-ML) using FastTree-2 on 51-taxon datasets that evolved under a moderate amount of ILS but with varying rates of HGT under the stochastic HGT model. In our analyses, all methods produced binary trees; hence, we report the normalized bipartition distance (also called the Robinson-Foulds [44] distance) between estimated species trees and true species trees. We report results for both true and estimated gene trees, with 10 to 1000 genes. To evaluate the relationship between topological accuracy and performance with respect to the optimization problem that ASTRAL-2 and wQMC attempt to solve, we compared the quartet support scores and topological accuracy of trees computed by ASTRAL-2 and wQMC.

### Results on estimated gene trees

For datasets with 10 genes (Fig. 2), all the methods are very similar when there is no HGT (i.e., HGT rate (1)), with error rates varying from 13.0% (ASTRAL-2 and wQMC) to 14.5% (NJst). Error rates increase with increasing HGT rates, but the increases are generally small until HGT rate (4), where all methods have error between 14.9% (ASTRAL-2) and 16.8% (CA-ML). Furthermore, the differences between methods remain small (no more than 1.9% between the methods) through HGT rate (4). However, there are substantial differences between methods under the two highest HGT rates (5) and (6), with CA-ML having the highest error (26.6% and 40.2%, respectively) and ASTRAL-2 having the least error (18.4% and 28.1%, respectively). While the differences between wQMC and NJst were often small, typically wQMC was more accurate than NJst.

**Figure 2.**
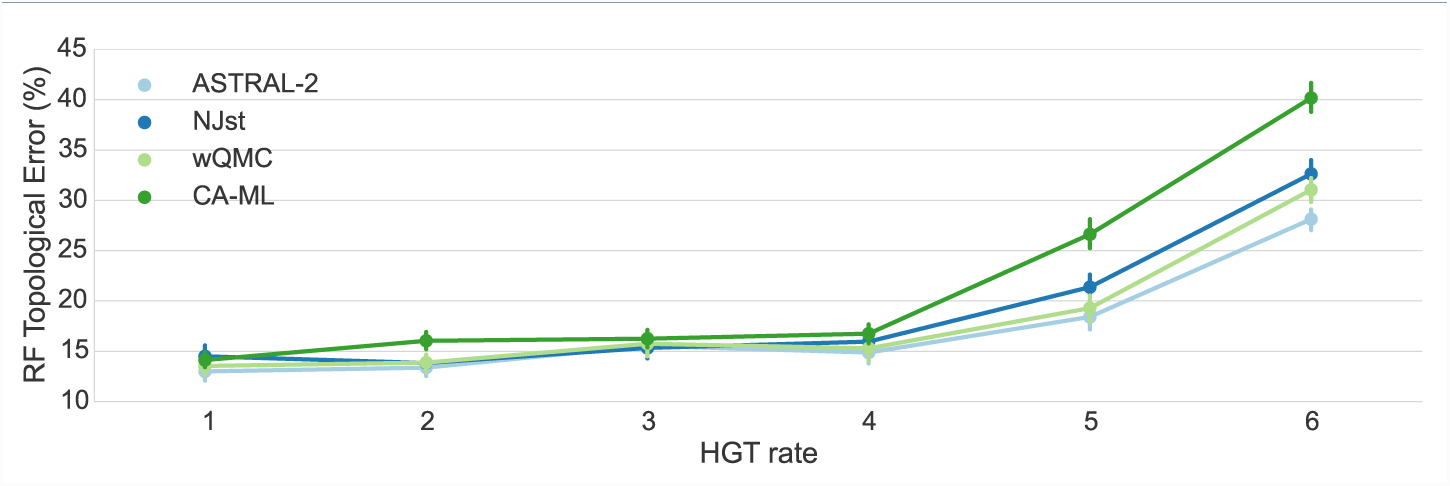
Mean Robinson-Foulds error rate on datasets with 10 genes. We show mean RF error rates for summary methods applied to estimated gene trees as well as for an unpartitioned maximum likelihood concatenation analysis. Error bars indicate standard error; 50 replicates per dataset.

The same trends hold on datasets with larger numbers of genes (Fig. 3); in particular, ASTRAL-2 remains typically the most accurate method (or close to the most accurate method) and CA-ML is typically the least accurate. However, as the number of genes increase, the species tree estimation error drops for all methods, and the differences between methods become even smaller. For example, on 50 genes the maximum error for HGT rates (1)-(4) is 7.8% (CA-ML) and the smallest error is 7.3% (ASTRAL-2 and NJst). By 200 genes, the maximum error of all methods on HGT rates (1)-(4)) is 5.1% (NJst) and the smallest is 4.5% (ASTRAL-2). With 1000 genes, the maximum error on HGT rates (1)-(4) is only 3.1% (wQMC and NJst) and the lowest is 2.5% (CA-ML). However, under the two higher HGT rates (HGT rates (5) and (6)), the differences between methods can be noteworthy, even with large numbers of genes. More importantly, under these higher HGT rates, CA-ML is substantially less accurate than all of the summary methods. As an example, under HGT rate (6), CA-ML has 16.8% error on 50 genes, while ASTRAL-2 has 10.3% error. One interesting trend that is hard to explain is that error rates do not always increase with increases in HGT rates; for example, results on 1000 estimated trees show some small decrease in error for ASTRAL-2 and NJst between HGT rates (4) and (6). Finally, while ASTRAL-2 is the most accurate of the summary methods, but the difference between ASTRAL-2 and the other summary methods is small (ranging from 0.3% to 1.9%). Indeed, the differences between the summary methods given 400 or more genes are very small — at most 0.9%.

**Figure 3.**
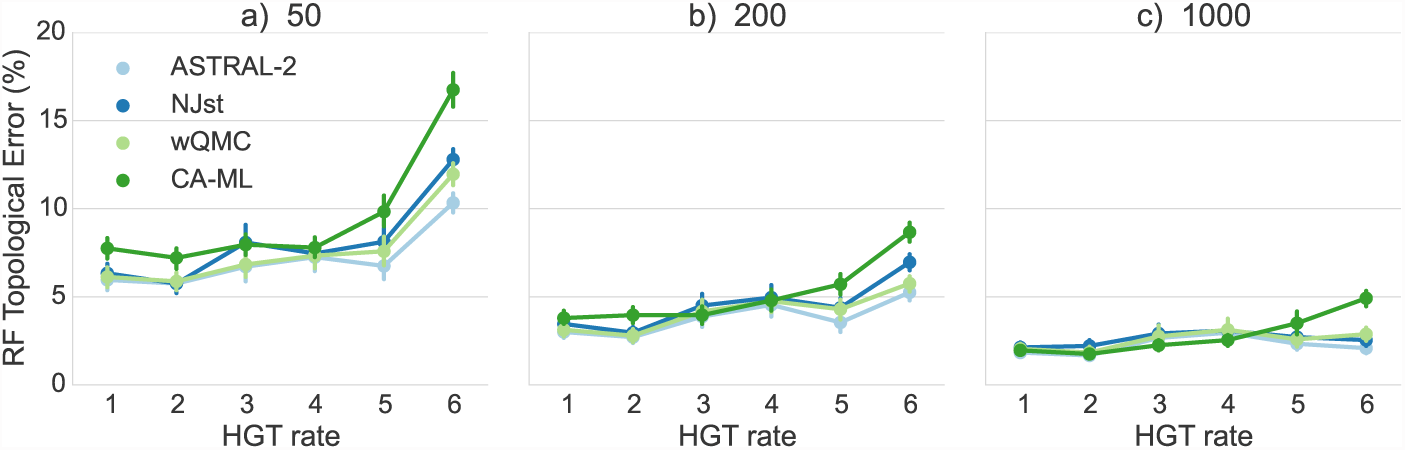
Mean Robinson-Foulds error rates on datasets with 50, 200, and 1000 estimated gene trees. We show results for summary methods applied to estimated gene trees as well as for an unpartitioned maximum likelihood concatenation analysis. Error bars indicate standard error; 50 replicates per dataset.

### Results on true gene trees

We show results on true gene trees in Figures 4 and 5. Unsurprisingly, error rates of species trees estimated on true gene trees are lower than those estimated on estimated gene trees; while the reduction depended on the model condition, for the ASTRAL-2 datasets with 1000 genes and HGT rate (1), we see a reduction of more than 50%. Differences between methods were reduced on the true gene trees, but otherwise, all the trends are the same as for estimated gene trees.

**Figure 4.**
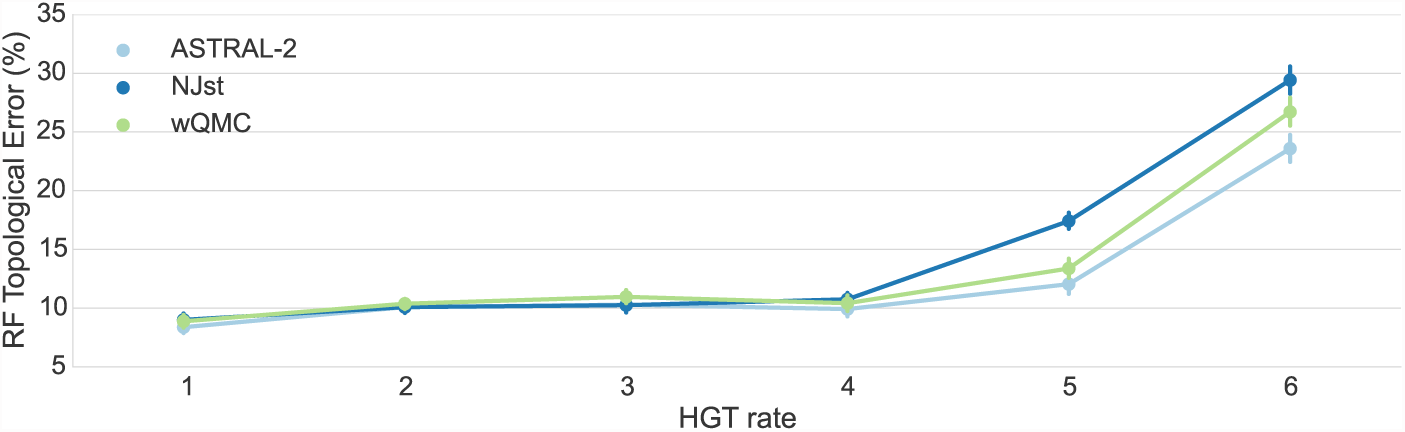
Mean Robinson-Foulds error rates on 10 true gene trees. We show mean RF error rates of summary methods applied to true gene trees; error bars indicate standard error. 50 replicates per model condition.

**Figure 5.**
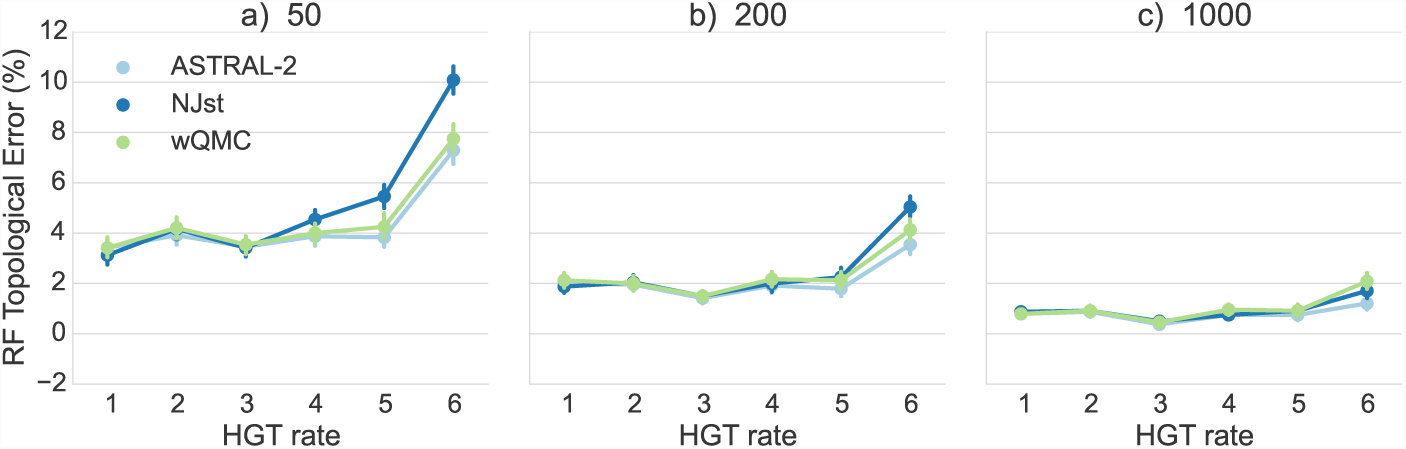
Mean Robinson-Foulds error rates on 50, 200, and 1000 true gene trees. We show mean RF error rates of summary methods applied to true gene trees; error bars indicate standard error. 50 replicates per model condition.

### Comparing quartet scores of trees produced by ASTRAL-2 and wQMC

While the differences between ASTRAL-2 and wQMC are often small, ASTRAL-2 nearly always matches or improves on wQMC with respect to tree topology. Both ASTRAL-2 and wQMC attempt to solve the Maximum Quartet Support Species Tree problem (MQSST, see Methods), but use very different techniques. In particular, ASTRAL-2 constrains the search space based on the input gene trees, and then finds an optimal solution within that constrained space, but wQMC uses a greedy heuristic and does not constrain the search. One hypothesis for the improved topological accuracy of ASTRAL-2 compared to wQMC is that ASTRAL-2 finds better solutions to the MQSST optimization problem, and a competing hypothesis is that the higher topological accuracy achieved by ASTRAL-2 is due in part to the constraint it imposes on the solution space.

We examined the quartet scores for wQMC and ASTRAL-2 across the different model conditions. For 57.2% of all cases involving estimated gene trees, the species trees returned by the two methods had the same quartet support. ASTRAL-2 returned a tree with a better quartet score than wQMC 29.8% of the time while wQMC returned a tree with a better quartet score 13.0% of the time. Thus, in general ASTRAL-2 does a better job than wQMC of finding good solutions to MQSST. However, there are cases in which wQMC produces trees with better scores, and the cases are typically cases with high HGT levels (i.e., there are no cases with HGT rate (1), and more than half of the cases occurred for HGT rate (6)).

We investigated the 29 replicates for which wQMC has a better quartet support score, and therefore does a better job of solving the MQSST problem (Fig. 6). ASTRAL-2 and wQMC had the same topological accuracy on 8 datasets, ASTRAL-2 was more topologically accurate on 12, and wQMC was more topologically accurate on 9. Thus, even for those cases where wQMC finds trees with better quartet support scores, ASTRAL-2 tends to match wQMC with respect to accuracy, or produce topologically more accurate trees. Since wQMC does not constrain the search space, this means that wQMC can find trees with better quartet scores but which are outside the constrained search space, and that constraining the search space seems to be beneficial with respect to topological accuracy. In other words, although ASTRAL-2 generally is a better heuristic for the MQSST problem, part of the reason it is more topologically accurate is due to the constraint it imposes on the search space.

**Figure 6.**
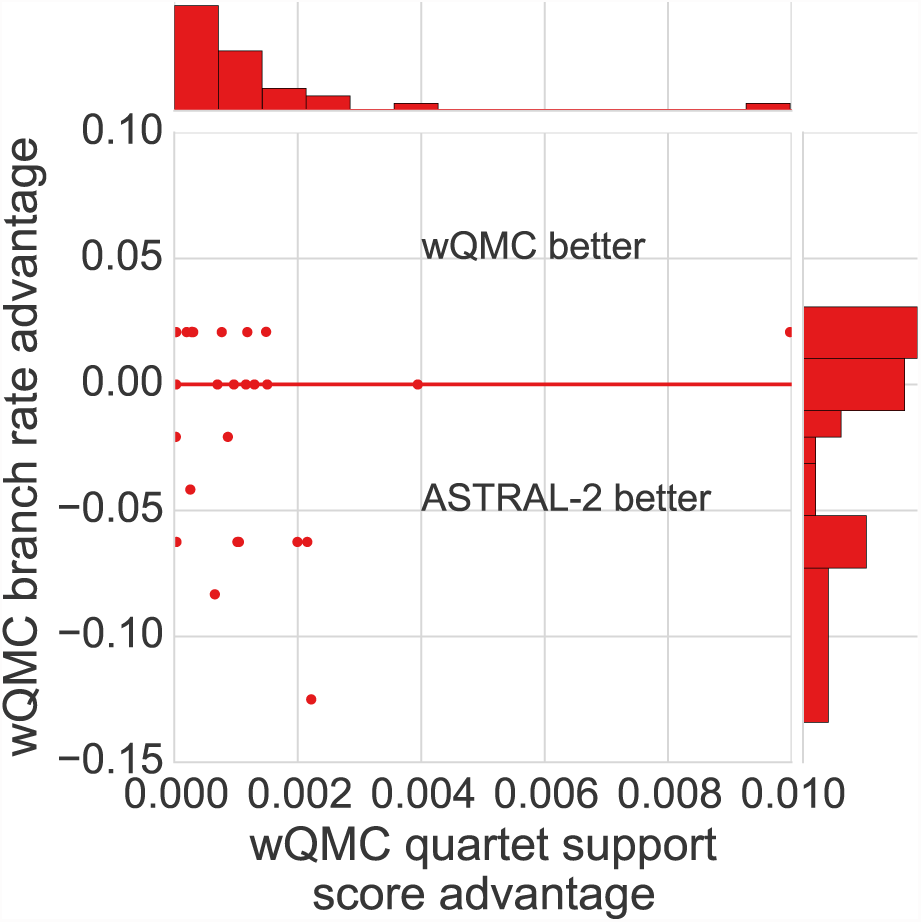
Scatterplot of differences in quartet support scores and topological error of wQMC and ASTRAL-2 trees. Each point (*x, y*) represents a dataset in which wQMC produced a tree with quartet support score *x* points higher than produced by ASTRAL-2, and with tree topological error *y* points lower. All values of *x* are strictly positive (we are only showing cases where wQMC produces a better quartet support score than ASTRAL-2), but values of *y* can be arbitrary. Points with *y <* 0 indicate datasets where ASTRAL-2 produces a topologically more accurate tree than wQMC, points with *y* = 0 indicate datasets where ASTRAL-2 and wQMC produce trees of equal accuracy, and points with *y >* 0 indicate datasets where ASTRAL-2 produces a tree that is topologically less accurate than wQMC. Of the points that are not on the *y* = 0 line, more are below the *y* = 0 line than above (i.e., 12 below compared to 9 above), indicating that ASTRAL-2 tends to produce more accurate tree topologies than wQMC on these datasets. Also, when wQMC is more accurate, the improvement is lower than when ASTRAL-2 is more accurate. Thus, even when wQMC finds trees with better quartet scores, ASTRAL-2 tends to produce more topologically accurate trees. Plots in the margins are histograms of the *x*– and *y*– axes.

### Cyanobacterial Data

We analyzed a cyanobacterial data set from [42] using ASTRAL-2 with multi-locus bootstrapping (see Methods) to estimate a species tree. Two estimated species trees were reported in [42]: one is the “plurality tree”, which has served as the reference tree for this dataset. The plurality tree is a supertree (computed using MRP [45]) on a set of quartet trees represented in a plurality of the gene trees that have high support. The other tree is a PhyML [46] maximum likelihood tree. The ASTRAL-2 majority consensus tree (see Methods) has 100% bootstrap support on all its branches, and is identical to the plurality tree; that has served as the reference tree for this dataset. The wQMC tree was previously reported for this dataset in [40], and is also topologically identical to the plurality tree.

## Discussion

While all methods had very good accuracy on the simulated datasets under the lowest HGT rates, they were clearly differentiated on the higher HGT rates, especially when the number of genes was not too large. Specifically, on the higher HGT rates, concatenation using maximum likelihood and NJst were both less accurate than ASTRAL-2 and wQMC. However, all summary methods we explored were impacted by gene tree estimation error. Furthermore, there are no proofs of convergence to the true species tree if the gene trees have estimation error for these or other standard summary methods [47, 12]. Since many of the lower HGT model conditions had substantial gene tree heterogeneity resulting from ILS, this study shows that many methods — and even unpartitioned concatenation using maximum likelihood - can be highly accurate under these highly heterogeneous model conditions.

Results on the biological dataset showed that ASTRAL-2 and wQMC both matched the reference “plurality tree”, and hence may be correct. But this analysis is perhaps less helpful, since the reference tree is based on the MRP analysis of a set of quartet trees, and MRP on quartet trees is a heuristic for the unweighted version of the optimization problem addressed by wQMC and ASTRAL-2. Thus, the three methods are closely related in terms of their optimality criteria, and this may explain why they produce the same tree on this input.

This experimental study evaluated the performance of these methods when HGT is also present, and demonstrated that wQMC and ASTRAL-2 maintained good accuracy even in the presence of HGT, while NJst tended to be more impacted by high levels of HGT. The explanation as to why NJst is not as robust to high HGT levels as ASTRAL-2 and wQMC is likely to be that the theoretical justification for NJst only applies to the MSC model, and not to the bounded HGT models. On the other hand, both ASTRAL-2 and wQMC attempt to solve the MQSST problem, for which optimal solutions are statistically consistent under the MSC model, and also under the bounded HGT models discussed in [36].

Finally, the slight advantage ASTRAL-2 had over wQMC in terms of topological accuracy is largely due to its better ability to find good solutions to the MQSST problem, but constraining the search space is also part of the reason that ASTRAL-2 has good topological accuracy, even under conditions with very high rates of HGT.

## Conclusions

This study evaluated ASTRAL-2, NJst, wQMC, and concatenated analysis using unpartitioned maximum likelihood (CA-ML) on one biological and several simulated datasets in which ILS and HGT were both present. We observed that the quartet-based methods (ASTRAL-2 and wQMC) generally had better accuracy than NJst, and that CA-ML could be more accurate than all methods under conditions with low HGT rates. In particular, ASTRAL-2, a species tree estimation method that was initially designed to estimate species trees in the presence of ILS, had excellent accuracy and generally gave somewhat more accurate results than the other methods we explored. However, all methods were highly accurate under the low to moderate HGT levels, and were only differentiated under the two highest HGT levels. The methods based on quartets (i.e., wQMC and ASTRAL-2) had the highest robustness to HGT. While the study is limited in scope, the results suggest that highly accurate species trees can be constructed, even in the presence of both HGT and ILS, using quartet-based methods.

As noted, ASTRAL-2 and NJst are statistically consistent under the MSC model (in which only ILS occurs), and ASTRAL-2 is also statistically consistent under the bounded HGT models addressed by [36]. However, NJst has not been shown to be statistically consistent under the bounded HGT models, and wQMC may not be statistically consistent under either model (because it is not guaranteed to solve its optimization problem exactly, even when all the dominant quartet trees are compatible). Because the proof of statistical consistency for ASTRAL-2 depends only on the requirement that for all sets of four taxa, the most probable quartet tree is topologically identical to the induced species tree on the four taxa, we conjecture that ASTRAL-2 will be statistically consistent under models in which both ILS and HGT occur but at bounded rates (where the bounds on one process will depend on the other’s bounds).

Although the results in this study are encouraging, future work needs to evaluate the performance of species tree estimation methods under a broader set of conditions. In particular, we only evaluated performance under the stochastic HGT model; future work should evaluate methods under the highways model as well. Our datasets had only one level of ILS, and it is possible that under conditions with higher or lower levels of ILS, the effect of HGT would be different. This study was limited to gene trees in which heterogeneity was due only to ILS and HGT; future studies should examine other sources of discord, including gene duplication and loss, and/or orthology detection errors. Larger numbers of taxa, and/or gene trees with missing taxa, are also likely to present significant analytical challenges, and accurate estimation may not be as easily obtained. Hence, future studies should also evaluate accuracy on larger and more challenging datasets, in order to determine whether the good accuracy we saw for the quartet-based methods is maintained under more difficult conditions. Similarly, it is possible that some methods might provide highly accurate results on smaller numbers of species, and that the relative performance of methods could change on those conditions. Thus, performance on small datasets (with perhaps only 10 species) should also be explored.

This study was limited in terms of the methods that were explored, in that we restricted the analysis to reasonably fast methods, and of these fast methods we only explored those methods that had been shown to perform well under ILS-only scenarios. However, it is possible that some coalescent-based species tree estimation methods, such as MP-EST, STAR, etc., might perform well under HGT+ILS scenarios. It is also likely some computationally intensive methods, such as BUCKy-pop, *BEAST, and BEST, might provide better accuracy than ASTRAL-2 on datasets with HGT+ILS. There are also methods designed to infer species trees in the presence of gene tree discordance resulting from duplication and loss, and it is possible that some of these methods (e.g., PhylDog [48] and MixTreEM [49]) might have good accuracy under the MSC. Future work should also explore CA-ML using different ML heuristics (e.g., PhyML [46], nhPhyML [50], IQTree [51]) and under more complex sequence evolution models. In addition, it would be very interesting to explore fully partitioned ML analyses, since these have very different statistical properties than unpartitioned analyses [12].

## Methods

Species tree estimation methods

*Maximum Quartet Support Species Tree Problem*

ASTRAL, ASTRAL-2, and wQMC all address the same optimization problem, which we now explain. Given an input set *G* of gene trees on a species set *S* and a quartet tree *q* on four species from *S*, we let *n*(*G, q*) denote the number of gene trees in *G* that induce the quartet tree *q*. Then, the *quartet support* of *T* given *G*, denoted *w*_G_ (*T*), is ∑_q∈Q(T)_ *n*(*G, q*), where Q(*T*) denotes the set of all quartet trees in *T*. Hence, we can define the *Maximum Quartet Support Species Tree Problem* (MQSST), as follows.

- Input: a set of gene trees *G* on a species set *S*.
- Output: a tree *T* on the species set *S* maximizing *w*_G_ (*T*), the quartet support of *T* given *G*.

MQSST is *N P*-hard when the input set of gene trees induce only one tree for each set of four taxa in *S* [52], and is of unknown computational complexity when all the gene trees are complete (i.e., have all the species in *S*).

### Weighted Quartets MaxCut

The quartet amalgamation method wQMC [40] is a greedy heuristic for a weighted version of the MQSST problem, in which the input can have weights on each quartet tree. The wQMC heuristic uses a greedy strategy to find good solutions to its optimization problems, but is not guaranteed to solve its optimization problem (weighted MQSST) exactly. To use wQMC as a summary method, we define the weight of a quartet tree *q* to be the quartet support *n*(*G, q*) of *q* in the input set of gene trees *G*.

We wrote scripts (available in our supporting online material) that use a previously published code [53] to compute the weights of each quartet tree. After we calculate these weights (saving them in a file called `<quartetscores>`), we run wQMC version

3.0 using the following command:

~~~
./max-cut-tree qrtt=<quartetscores> weights=on otre=<speciestree>
~~~

### ASTRAL and ASTRAL-2

ASTRAL [18] and its improved version, ASTRAL-2 [19], also attempt to solve the MQSST problem. Both have exact versions that provably solve the MQSST problem but run in exponential time, and faster versions that constrain the search space (using the input set of gene trees), and then provably solve the constrained problem exactly. ASTRAL and ASTRAL-2 differ in how they constrain the search space (ASTRAL-2 searches a larger part of tree space than ASTRAL) and how they are implemented (ASTRAL-2 is faster). Here we focus on ASTRAL-2, since it is faster and more accurate than ASTRAL.

Given the input set of gene trees, ASTRAL-2 defines a set *χ* of bipartitions on the taxon set *S*; when all the gene trees are complete (i.e., have no missing taxa), then *χ* will contain all the bipartitions from the input gene trees as well as potentially other bipartitions. ASTRAL-2 runs in *O*(*nk*|*χ* |^2^) time, where *n* is the number of species and *k* is the number of genes, and thus can be fast whenever |*χ* | is not too large. While |*χ* | is not theoretically bounded by a polynomial in *n* and *k*, for many datasets |*χ* | is not very large, so that ASTRAL-2 is able to complete analyses within 24 hours on 1000 species and 1000 genes [19].

ASTRAL-2 finds a globally optimal solution to the constrained optimization problem where we restrict the output species tree to draw its bipartitions from χ. ASTRAL and ASTRAL-2, run in their default versions (which use the constrained search), are both statistically consistent under the multispecies coalescent model when all the gene trees are complete (i.e., this restriction to the set χ of bipartitions does not change their statistical guarantees) [19].

We now provide a proof for Theorem 3, establishing that ASTRAL and ASTRAL2, run in default mode, are statistically consistent under the MSC model and also under the bounded HGT models.

**Proof for Theorem 3**. As proved in [18, 19], ASTRAL and ASTRAL-2 are guaranteed to find globally optimal solutions to the constrained MQSST problem. The default settings for the constraint set χ of bipartitions allowed in the output species tree always includes all bipartitions from the input gene trees; hence, as the number of genes increases, with probability converging to 1, every bipartition from the species tree will be in the set χ. Therefore, with probability converging to 1, the true species tree will be a feasible solution (i.e., within the constrained search space) as the number of loci and number of sites per locus both increase (as established in [18, 19]). Recall that the quartet support score of a tree *T* is the total, over all quartet trees in *T*, of the number of gene trees that contain that quartet tree. As shown in [36], under the bounded HGT models in [36], the most probable quartet tree on any four taxon set *A* is topologically identical to the quartet tree on *χ* induced by the true species tree. Hence, with probability converging to 1, under these bounded HGT models, the most frequent quartet tree on any set *A* of four leaves will be the true species tree on *A*. Given any set of gene trees in which for all four-leaf sets *A* the most frequent quartet tree on *A* is the true species tree on *A*, the quartet support score of the true species tree *T* ^*^ will be the maximum possible quartet support score (since any other species tree *T* cannot have larger quartet support for any quartet tree). Furthermore, given any set of gene trees in which the most frequent quartet tree is unique for all four taxa and equal to the species tree on the four taxa, the true species tree *T* ^*^ will have the unique maximum quartet support score. Hence, as the number of loci and number of sites per locus both increase, the tree returned by an exact solution to the constrained MQSST problem, using default settings for χ, will converge in probability to the true species tree *T* ^*^. Therefore, ASTRAL and ASTRAL-2 are statistically consistent under the bounded HGT models of [36].

We ran ASTRAL-2 version 4.7.6 on the simulated data using the following command:

~~~
java -jar astral.4.7.6.jar -i <genetrees> -o <speciestree>
~~~

where `<genetrees>` is a file containing the gene trees in newick format, and `<speciestree>` is the output.

For the biological data, we used ASTRAL-2 with multi-locus bootstrapping (MLBS), using the following commands:

~~~
java -jar astral.4.7.6.jar -i < bootstrap replicates >
-o <species replicate>
~~~

where bootstrap replicates is the collection of 1128 gene trees generated by taking the *n*^th^ line of the gene tree file *n* = {1, …, 100}, and species replicate is the *n*^th^ bootstrap replicate species tree *T*_n_. To calculate the final species tree *T* with bootstrap support values, we computed the majority consensus tree using Dendropy version 3.12.2 [54].

### NJst

NJst is a summary method that has two steps. In the first step, it computes a distance matrix on the species set, where *D*[*x, y*] is the average leaf-to-leaf topological distance between *x* and *y* among all the gene trees. In the second step, it runs neighbor joining [55], a popular distance-based phylogeny estimation method. NJst is statistically consistent under the MSC model because the distance matrix it computes converges in probability to an additive matrix defining the true species tree, and neighbor joining will return the true species tree once the computed distance matrix is sufficiently close to the additive matrix for the species tree; see [17] for this proof.

To run NJst, we used phybase version 1.4 [56] and custom scripts, available in our supplementary material.

### Gene tree estimation

To compute gene trees, we ran FastTree-2 version 2.1.4, using the following command:

~~~
fasttree -nt -gtr -quiet -nopr -gamma -n 1000 [input] > [output]
~~~

 where `[input]` is a file that includes all the alignments of all 1000 genes and `[output]` will be one file with all 1000 estimated gene trees.

## CA-ML

To perform the concatenated analyses under maximum likelihood, we ran FastTree2 version 2.1.4, with the following command:

~~~
fasttree -nt -gtr -nopr [input] > [output]
~~~

## Computing Error Rates

The coalescent-based methods ASTRAL-2, wQMC, and NJst used in this study all return binary species trees. We also verified that all trees returned in our CA-ML analysis were binary, and all simulated data used in this study contained only binary model species trees. The Robinson-Foulds (RF) distance [44] between two trees *T*_1_ and *T*_2_ on the same set of *n* taxa measures the number of bipartitions that appear in only one of *T*_1_ or *T*_2_. Therefore, if *T*_1_ and *T*_2_ are identical, the RF distance is 0, and the maximum RF distance between *T*_1_ and *T*_2_ is 2*n*-6. The RF distance can be converted to an error rate by dividing by 2*n* 6. When comparing only binary trees, false negative rates, false positive rates, and normalized Robinson-Foulds distances are all equivalent. Therefore, we computed missing branch rates to establish error rates, but we report RF rates. Error rates were computed by finding the missing branch rate using custom scripts available in our supporting online materials.

## Measuring Quartet Support Scores of ASTRAL-2 and wQMC

The command used to measure the quartet support score was

~~~
java -jar astral.4.7.6.jar -q <speciestreefile> -i <genetreesfile>
~~~

## Data

*HGT+ILS Simulated Data* The simulated dataset was simulated using SimPhy

[57] version 1.0 (downloaded January 20, 2015). There are 6 data sets containing 50 replicates apiece: each replicate has its own 51-taxon species tree. For every model species tree, one taxon is an outgroup, and so is actually a 50-taxon rooted species tree. These model trees were simulated under a Yule process, with birth rates set to 0.000001 (per generation) and the maximum tree length set to 2 million generations. Then, on each species tree, 1000 locus trees are simulated, where each can differ from the species tree due to HGT events, and we used HGT rates (1)-(6) given by 0, 2 × 10^-9^, 5 × 10^-9^, 2 × 10^-8^, 2 × 10^-7^, and 5 × 10^-7^. These values correspond to expected numbers of HGT events per gene of 0, 0.08, 0.2, 0.8, 8, and 20. Thus, HGT rate (1) is no HGT events, HGT rate (2) is 0.08 HGT events per gene, up to HGT rate (6) of 20 HGT events per gene. Note that in our simulations, for each HGT event, the probability of a branch being chosen as the receptor of the transfer is proportional to its distance from the donor.

Once locus trees are simulated, a gene tree is simulated for each locus tree according to the MSC model, with population size parameter set to 200,000. Thus, at the end, we have 1000 true genes that differ from the species tree due to both ILS and also potentially HGT (when the HGT rate is positive).

The SimPhy command used to generate a model replicate in the data sets is

~~~
simphy -rs 50 -rl U:1000,1000 -rg 1 -st U:2000000,2000000 -si U:1,1
-sl U:50,50 -sb U:0.000001,0.000001 -cp U:200000,2000000
-hs L:1.5,1 -hl L:1.2,1 -hg l:1.4,1 -cu E:10000000 -so U:1,1 -od 1
-or 0 -v 3 -cs 293745 -o model.50.2000000.0.000001.<transferrate>
-lt U:<transferrate>,<transferrate> -lk 1
~~~

On each simulated true gene tree, we used INDELible [58] v. 1.03 to simulate sequence alignments according to the GTR+Γ model, with model parameters estimated from three different real datasets (these parameters are identical to those used in [19]). This simulation produces GTR parameters that vary from one gene to another, where the parameters are drawn for each gene from a distribution at random. See [19] for details about the simulation process. The alignment length is set to 1000bp for all genes. After simulating gene alignments, we used FastTree-2 [41] to estimate gene trees under the GTR model. Thus for each replicate, we have both true and estimated gene trees.

For HGT rate (1) (where all the discordance is due to ILS), the average RF distance between true gene trees and the species tree is 30.4%. Therefore, the amount of ILS in these data sets is moderately high.

*Cyanobacterial Data* The cyanobacterial data set has 1128 genes on 11 taxa, and was first analyzed in [42], which suggested that the 11 genome sequences may have acquired between 9.5% and 16.6% of their genes through HGT. We obtained 100 bootstrap replicate gene trees for each of the 1128 genes from the first author of [43], and computed an ASTRAL-2 tree on these data using multi-locus bootstrapping.

## Availability of supporting data

All data used in this study, and commands needed to regenerate the data, are available online at goo.gl/0p4IGD.

## Abbreviations

CA-ML: concatenated analysis using maximum likelihood
GTR+Gamma: Generalized Time Reversible model of site evolution with Gamma distributed rates across sites HGT: horizontal gene transfer
HGT: horizontal gene transfer
ILS: incomplete lineage sorting
MSC: multi-species coalescent
ML: maximum likelihood
MLBS: multi-locus bootstrapping
MQSST: maximum quartet support species tree
MSC: multi-species coalescent

## Competing interests

The authors declare that they have no competing interests.

## Author’s contributions

RD performed ASTRAL-2 analyses of the simulated and biological data sets, the CA-ML on the simulated data for 10 genes, and wrote the first draft of the paper. PV performed the wQMC and NJst analyses of the simulated data sets and made figures. SM generated the simulated data, performed the CA-ML analysis for 50, 200, and 1000 genes, and made figures. TW conceived of the project, supervised the research, proved the theorems, and wrote the final paper. All authors read and critiqued drafts of the paper.

## Acknowledgements

RD was supported by NSF grant DMS-1401591. PV was supported by the Roy J. Carver graduate fellowship from the UIUC College of Engineering. SM was supported by a Howard Hughes Medical Institute graduate fellowship and by NSF grant DBI-1461364. TW was supported by NSF grant DBI-1461364 and by a gift from the Grainger Foundation to the University of Illinois at Urbana-Champaign College of Engineering. The authors thank Mukul Bansal for sharing the data from [42].

## References

1. Morrison, D.A.: Introduction to Phylogenetic Networks. RJR Productions, Uppsala, Sweden (2011)

2. Sjölander, K.: Phylogenomic inference of protein molecular function: advances and challenges. Bioinformatics 20(2), 170–179 (2004)

3. Eisen, J.A., Fraser, C.M.: Phylogenomics: intersection of evolution and genomics. Science 300(5626), 1706–1707 (2003)

4. Engelhardt, B.E., Jordan, M.I., Repo, S.T., Brenner, S.E.: Phylogenetic molecular function annotation. J Phys: Conf Ser 180, 12024 (2009)

5. Thompson, J.N.: The Geographic Mosaic of Coevolution. The University of Chicago Press, Chicago (2005)

6. Alberts, B., Johnson, A., Lewis, J., Raff, M., Roberts, K., Walte, P.: Molecular Biology of the Cell, 4th edn. Garland Science, New York (2002)

7. Nussbaum, R., McInnes, R.R., Willard, H.F.: Genetics in Medicine, 7th edn. Saunders Elsevier, Philadelphia, PA (2007)

8. Arnold, M.L.: Natural Hybridization and Evolution. Oxford University Press, Oxford (1997)

9. Maddison, W.: Gene trees in species trees. Syst Biol 46, 523–536 (1997)

10. Woese, C.: On the evolution of cells. Proc Natl Acad Sci USA 99, 8742–8747 (2002)

11. Kingman, J.F.C.: On the genealogy of large populations. J Appl Probab 19A, 27–43 (1982)

12. Warnow, T.: Concatenation analyses in the presence of incomplete lineage sorting. PLOS Currents: Tree of Life 105, 10–13718410131445951717 (2015)

13. Liu, L., Yu, L., Edwards, S.V.: A maximum pseudo-likelihood approach for estimating species trees under the coalescent model. BMC Evol Biol 10(1), 302 (2010)

14. Mossel, E., Roch, S.: Incomplete lineage sorting: consistent phylogeny estimation from multiple loci. IEEE/ACM Trans Comput Biol Bioinformatics (TCBB) 7(1), 166–171 (2011)

15. Kubatko, L.S., Carstens, B.C., Knowles, L.L.: STEM: species tree estimation using maximum likelihood for gene trees under coalescence. Bioinformatics 25(7), 971–973 (2009)

16. Liu, L., Yu, L., Pearl, D.K., Edwards, S.V.: Estimating species phylogenies using coalescence times among sequences. Syst Biol 58(5), 468–477 (2009)

17. Liu, L., Yu, L.: Estimating species trees from unrooted gene trees. Syst Biol 60, 661–667 (2011)

18. Mirarab, S., Reaz, R., Bayzid, M.S., Zimmerman, T., Swenson, M., Warnow, T.: ASTRAL: genome-scale coalescent-based species tree estimation. Bioinformatics 30, 1541–1548 (2014)

19. Mirarab, S., Warnow, T.: ASTRAL-II: coalescent-based species tree estimation with many hundreds of taxa and thousands of genes. Bioinformatics 31 (2015). doi:10.1093/bioinformatics/btv234

20. Jarvis, E.D., Mirarab, S., et al.: Whole genome analyses resolve early branches in the tree of life of modern birds. Science 346(6215), 1320–1331 (2014)

21. Wickett, N.J., Mirarab, S., Nguyen, N., Warnow, T., Carpenter, E., Matasci, N., Ayyampalayam, S., Barker, M.S., Burleigh, J.G., Gitzendanner, M.A., Ruhfel, B.R., Wafula, E., Der, J.P., Graham, S.W., Mathews, S., Melkonian, M., Soltis, D.E., Soltis, P.S., Miles, N.W., Rothfels, C.J., Pokorny, L., Shaw, A.J., DeGironimo, L., Stevenson, D.W., Surek, B., Villarreal, J.C., Roure, B., Philippe, H., dePamphilis, C.W., Chen, T., Deyholos, M.K., Baucom, R.S., Kutchan, T.M., Augustin, M.M., Wang, J., Zhang, Y., Tian, Z., Yan, Z., Wu, X., Sun, X., Wong, G.K.-S., Leebens-Mack, J.: Phylotranscriptomic analysis of the origin and early diversification of land plants. Proc Natl Acad Sci USA 111(45), 4859–4868 (2014). doi:10.1073/pnas.1323926111. http://www.pnas.org/content/111/45/E4859.full.pdf+html

22. Roch, S., Steel, M.: Likelihood-based tree reconstruction on a concatenation of aligned sequence data sets can be statistically inconsistent. Theoret Popul Biol 100, 56–62 (2015)

23. Gatesy, J., Springer, M.S.: Concatenation versus coalescence versus “concatalescence“. Proc Natl Acad Sci USA 110 (2013). doi:10.1073/Proc. Natl. Acad. Sci.1221121110

24. Patel, S., Kimball, R., Braun, E.: Error in phylogenetic estimation for bushes in the tree of life. J Phylogen Evol Biol 1(110), 2 (2013)

25. Bayzid, M.S., Warnow, T.: Naive binning improves phylogenomic analyses. Bioinformatics 28, 2277–2284 (2013)

26. Liu, L.: BEST: Bayesian estimation of species trees under the coalescent model. Bioinformatics 24(21), 2542–2543 (2008)

27. Heled, J., Drummond, A.J.: Bayesian inference of species trees from multilocus data. Mol Biol Evol 27(3), 570–580 (2010)

28. Larget, B.R., Kotha, S.K., Dewey, C.N., Ané, C.: BUCKy: gene tree/species tree reconciliation with Bayesian concordance analysis. Bioinformatics 26, 2910–2911 (2010)

29. Zimmermann, T., Mirarab, S., Warnow, T.: BBCA: Improving the scalability of *BEAST using random binning. BMC Genomics 15(Suppl 6), 11 (2014)

30. Yang, J., Warnow, T.: Fast and accurate methods for phylogenomic analyses. BMC Bioinformatics 12, 4 (2011)

31. Chifman, J., Kubatko, L.: Quartet inference from SNP data under the coalescent model. Bioinformatics, 530 (2014)

32. Allman, E.S., Degnan, J.H., Rhodes, J.A.: Identifying the rooted species tree from the distribution of unrooted gene trees under the coalescent. J Math Biol 62, 833–862 (2011)

33. Galtier, N.: A model of horizontal gene transfer and the bacterial phylogeny problem. Syst Biol 56, 633–642 (2007)

34. Beiko, R.G., Harlow, T.J., Ragan, M.A.: Highways of gene sharing in prokaryotes. Proc Natl Acad Sci USA 102, 14332–14337 (2005)

35. Steel, M., Linz, S., Huson, D.H., Sanderson, M.J.: Identifying a species tree subject to random lateral gene transfer. J Theor Biol 322, 81–93 (2013)

36. Roch, S., Snir, S.: Recovering the tree-like trend of evolution despite extensive lateral genetic transfer: A probabilistic analysis. J Comput Biol 20, 93–112 (2013)

37. Snir, S., Rao, S.: Quartets MaxCut: A fast algorithm for amalgamating quartet trees. Mol Phylog Evol 62, 1–8 (2012)

38. Reaz, R., Bayzid, M.S., Rahman, M.S.: Accurate phylogenetic tree reconstruction from quartets: A heuristic approach. PloS One 9(8), 104008 (2014)

39. Chung, Y., Ané, C.: Comparing two Bayesian methods for gene tree/species tree reconstruction: simulations with incomplete lineage sorting and horizontal gene transfer. Syst Biol 60, 261–275 (2011)

40. Avni, E., Cohen, R., Snir, S.: Weighted quartets phylogenetics. Syst Biol 64, 233–242 (2015)

41. Price, M.N., Dehal, P.S., Arkin, A.P.: FastTree 2: approximately maximum-likelihood trees for large alignments. PloS ONE 5, 9490 (2010)

42. Zhaxybayeva, O., Gogarten, J.P., Charlebois, R.L., Doolittle, W.F., Papke, R.T.: Phylogenetic analyses of cyanobacterial genomes: quantification of horizontal gene transfer events. Genome Res 16, 1099–1108 (2006)

43. Bansal, M.S., Banay, G., Gogarten, J.P., Harlow, T.J., Shamir, R.: Systematic inference of highways of horizontal gene transfer in prokaryotes. Bioinformatics 29, 571–579 (2013)

44. Robinson, D.F., Foulds, L.R.: Comparison of phylogenetic trees. Math Biosci 53, 131–147 (1981)

45. Baum, B.R., Ragan, M.A.: The MRP method. In: Bininda-Emonds, O.R.P. (ed.) Phylogenetic Supertrees: Combining Information to Reveal The Tree Of Life, pp. 17–34. Kluwer Academic, Dordrecht, the Netherlands (2004)

46. Guindon, S., Gascuel, O.: A simple, fast, and accurate algorithm to estimate large phylogenies by likelihood. Syst Biol 52, 696–704 (2003)

47. Roch, S., Warnow, T.: On the robustness to gene tree estimation error (or lack thereof) of coalescent-based species tree methods. Syst Biol, 10–1093016 (2015)

48. Boussau, B., Szölloősi, G.J., Duret, L., Gouy, M., Tannier, E., Daubin, V.: Genome-scale coestimation of species and gene trees. Genome Res 23(2), 323–330 (2013)

49. Ullah, L., Parviainen, P., Lagergren, J.: Species tree inference using a mixture model. Mol Biol Evol (2015). doi: 10.1093/molbev/msv115

50. Boussau, B., Gouy, M.: Efficient likelihood computations with non-reversible models of evolution. Syst Biol 55(5), 756–68 (2006)

51. Nguyen, L.-T., Schmidt, H.A., von Haeseler, A., Minh, B.Q.: IQ-TREE: A fast and effective stochastic algorithm for estimating maximum-likelihood phylogenies. Mol Biol Evol 32(1), 268–274 (2015)

52. Jiang, T., Kearney, P., Li, M.: A polynomial time approximation scheme for inferring evolutionary trees from quartet topologies and its application. SIAM J Comput 30, 1942–1961 (2001)

53. Johansen, J.: Computing triplet and quartet distances. PhD thesis, Aarhus University, Computer Science Department (2013)

54. Sukumaran, J., Holder, M.T.: DendroPy: A Python library for phylogenetic computing. Bioinformatics 26, 1569–1571 (2010)

55. Saitou, N., Nei, M.: The neighbor-joining method: a new method for reconstructing phylogenetic trees. Mol Biol Evol 4, 406–425 (1987)

56. Liu, L.:Phybase server. https://faculty.franklin.uga.edu/lliu/content/phybase

57. Mallo, D., Oliviera Martins, L., Posada, D.: SimPhy: Comprehensive simulation of gene, locus and species trees at the genome-wide level. Available online. https://code.google.com/p/simphy-project/

58. Fletcher, W., Yang, Z.: INDELible: a flexible simulator of biological sequence evolution. Mol Biol Evol 26, 1879–1888 (2009)

